# Neuroretinal-derived caveolin-1 promotes endotoxin-induced inflammation in the murine retina

**DOI:** 10.1101/2020.01.08.899377

**Authors:** Jami M. Gurley, Grzegorz Gmyrek, Mark E. McClellan, Stefanie M. Hauck, Mikhail G. Dozmorov, Jonathan D. Wren, Daniel J. J. Carr, Michael H. Elliott

## Abstract

Chronic ocular inflammation is associated with many retinal degenerative diseases that result in vision loss. The immune-privileged environment and complex organization of retinal tissue allow for the retina’s essential role in visual function processes, yet confound inquiries into cell-specific inflammatory effects that lead to retinal dysfunction and degeneration. Caveolin-1 (Cav1) is an integral membrane protein expressed in many retinal cell populations and has been implicated in retinal immune regulation. However, the direction (i.e., promotion or inhibition) in which Cav1 regulates inflammatory processes in the retina (as well as in other tissues) remains unclear. Previously, we showed that global-Cav1 depletion in the retina paradoxically resulted in reduced retinal inflammatory cytokine production with concurrent elevated retinal immune cell infiltration. We hypothesized that our previous results could be explained by cell-specific Cav1 functions in the retina. Here, we utilized our Chx10 (visual system homeobox 2)-Cre knockout model to deplete Cav1 specifically in the neural retinal (NR) compartment in order to clarify the role of neural retinal-specific Cav1 (NR-Cav1) in the retinal immune response to intravitreal LPS (lipopolysaccharide) challenge. Our data support that neural retinal-derived Cav1 *promotes* retinal tissue inflammation as Chx10-mediated Cav1 depletion was sufficient to suppress *both* retinal cytokine production and immune cell infiltration following inflammatory stimulation. Additionally, we identify Traf3 (tumor necrosis factor (TNF) receptor-associated factor 3) as a highly expressed potential immune modulator in retinal tissue that is upregulated with NR-Cav1 depletion. Furthermore, this study highlights the importance for understanding the role of Cav1 (and other proteins) in cell-specific contexts.

## INTRODUCTION

Chronic ocular inflammation promotes the development and progression of retinal degenerative diseases that lead to vision loss. Many of these retinal degenerative diseases, such as age-related macular degeneration, diabetic retinopathy, and glaucoma, are major causes of blindness and are strongly linked to dysregulated immune responses that damage retinal tissue and decrease visual function (1).

The retina, which is located in the back of the eye and is responsible for processing light in order to facilitate visual function, is a complex and delicate tissue that is highly susceptible to inflammation-mediated damage (1,2). As it is an immune-privileged tissue, the retina can elicit an immune response when necessary, but can also engage immune-suppressive mechanisms in order to control inflammatory responses that could lead to collateral damage (3,4). Under retinal degenerative disease conditions, uncontrolled inflammatory activation and/or dysregulation of ocular immune suppression are key contributors toward the development and progression of retinal disease and vision loss (1). Unfortunately, the leading treatment option for dysregulated retinal inflammation is localized corticosteroid administration, which increases the risk for ocular conditions that exacerbate vision loss (e.g., cataracts, glaucoma). Thus, understanding how to control inflammatory responses in the context of the immune-privileged retinal environment is important for developing novel therapeutic interventions that will preserve retinal and visual function.

Several studies have implicated Cav1 as an immune regulator in lung, brain, and myeloid-derived immune cells (5–7). In most cell types, Cav1 is integrated into specialized plasma membrane domains called caveolae, which are involved in a variety of cellular processes including clathrin-independent endocytosis, endothelial transcytosis, lipid transport, vascular tone and contraction, fibrosis, cell proliferation, monocyte/macrophage differentiation, and signaling pathway regulation (6,8–15). The role of Cav1 and caveolae in immune regulation is complex. Collectively, existing studies suggest that Cav1-dependent inflammatory promotion or inhibition can occur via various mechanisms that depend on cell-specific contexts. For example, in the lung, Cav1 has been shown to *promote* inflammation by inhibiting eNOS-dependent downregulation of proinflammatory cytokine production (6); whereas, in macrophages, Cav1 *inhibits* TLR4 (Toll-like receptor 4)-mediated inflammatory stimulation via caveolar sequestration, which prevents receptor complex formation and downstream signaling (10).

In the retina, Cav1 affects important visual processes such as phagolysosomal digestion of photoreceptor outer segments by the retinal pigment epithelium (RPE) (16), maintenance of retinal vascular blood-retinal-barrier integrity (17–19), and retinal neovascularization (20). We and others have also implicated Cav1 function specifically in retinal inflammation and immune response modulation (21–23). However, the specific role for Cav1 in retinal immunity is as perplexing as it has proven to be in extraocular tissue immune responses, and has only begun to be investigated (21). Previously, we showed that global-Cav1 deletion resulted in suppression of endotoxin-mediated retinal inflammatory cytokine production, which suggested that Cav1 *promotes* retinal inflammatory activation (21). Surprisingly, Cav1 deletion enhanced retinal immune cell infiltration, which paradoxically implied that Cav1 function *suppresses* the immune response. Because Cav1 is expressed throughout retinal tissue in several retinal cell populations, we reasoned that these paradoxical phenotypes could be explained by differential cell-specific Cav1 functions in the retina.

The aim of this study was to identify the Cav1-expressing retinal compartment responsible for *promotion* of endotoxin-induced inflammation. Here, we show that neural retinal-derived Cav1 depletion is sufficient to suppress inflammatory processes in the retina, which highlights the importance of the neural retinal compartment in immune activation. This study (in conjunction with ours and other previous work) contributes to our understanding of complex cell-specific Cav1 functions. Additionally, we identify Traf3 as a potential retinal immune modulator that deserves future study in regards to our search for novel therapeutic interventions aimed at suppressing the retinal immune response under conditions of chronic ocular inflammation.

## Results

### Neural retinal depletion of CAV1 protein in retinal tissue

Our previously published results presented seemingly contradictory data where global-Cav1 depletion resulted in suppression of retinal inflammatory cytokine production with a concurrent elevation in retinal immune cell infiltration (21). Because we hypothesized that this paradoxical result was due to divergent cell-specific Cav1 functions in different retinal cell populations, we wished to separate potential cell-specific Cav1 effects using conditional knockout animals. Thus, we utilized our previously-developed Chx10-Cre/Cav1^flox/flox^ neural retinal knockout animals (NR-Cav1 KO). This model expresses Cre recombinase in neuroretinal progenitors resulting in Cav1 gene recombination in retinal neurons and Müller glia but not other retinal glia (i.e., astrocytes or microglia) (Fig. 1A) (24). To validate specific targeting of retinal tissue, we first assessed CAV1 protein expression (via Western blot) in various tissues collected from NR-Cav1 wild type (WT) and KO animals (Fig. 1B-C). Whole retinal tissue lysates from NR-Cav1 KO animals displayed a ~76% depletion of CAV1 protein compared to their WT counterparts. Residual CAV1 protein expression is likely primarily due to the remaining CAV1 protein expression in non-targeted retinal cell populations (e.g., retinal vascular cells) with some modest contribution from mosaic recombination in the neural retina (Fig. 2) (34). There was no genotype-dependent difference in CAV1 protein expression in the other tissues we assessed (brain, lung, heart, spleen, or liver). Thus, our data show that NR-specific deletion targets a significant percentage of total retinal Cav1 expression.

**Figure 1.**
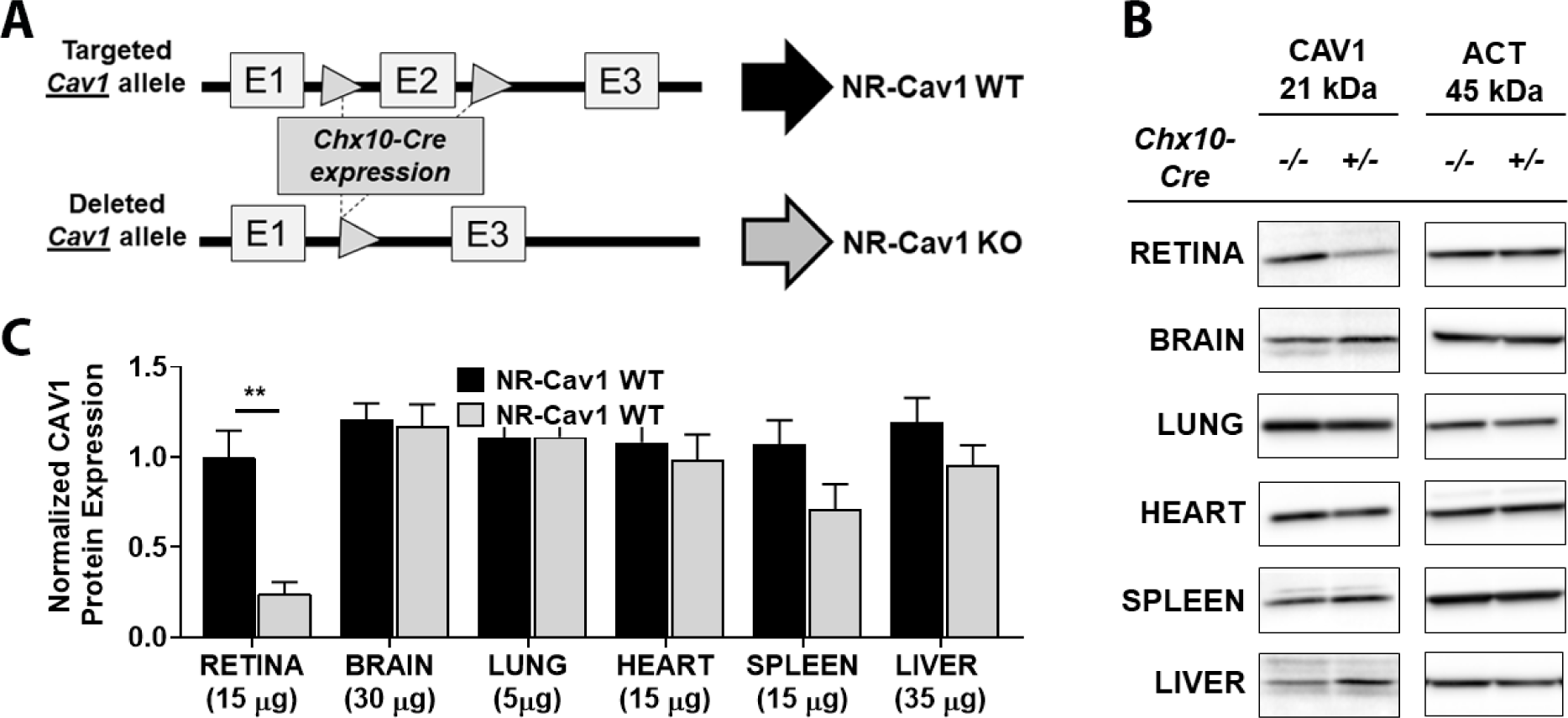
Effect of neuroretinal-specific deletion on whole retinal tissue CAV1 protein expression (Chx10-Cre/Cav1-flox model). (A) Schematic diagram illustrating the strategy for neuroretinal-specific *Cav1* deletion via Chx10-promoter driven Cre-mediated recombination of the floxed *Cav1* gene. (B) Representative Western blots of whole cell lysates from various tissues showing retina-specific CAV1 protein reduction following Chx10-Cre expression in the floxed *Cav1* model. Protein loads ranged from 5 µg to 35 µg, depending on tissue, in order to detect protein within the linear range of the assay. (C) Histogram of Western blot protein densitometry data quantification showing retinal tissue specific CAV1 protein reduction following NR-*Cav1*-depletion in Chx10-Cre/Cav1^+/+^ mice compared to Chx10-Cre/Cav^−/−^ controls. *β*-actin was used as a loading control and for normalization. Data are mean ± SEM and were analyzed via Student’s t-test (** p<0.01; N=7). NR, neural retina/neuroretinal; CAV1, Caveolin 1; ACT = β-actin.

**Figure 2.**
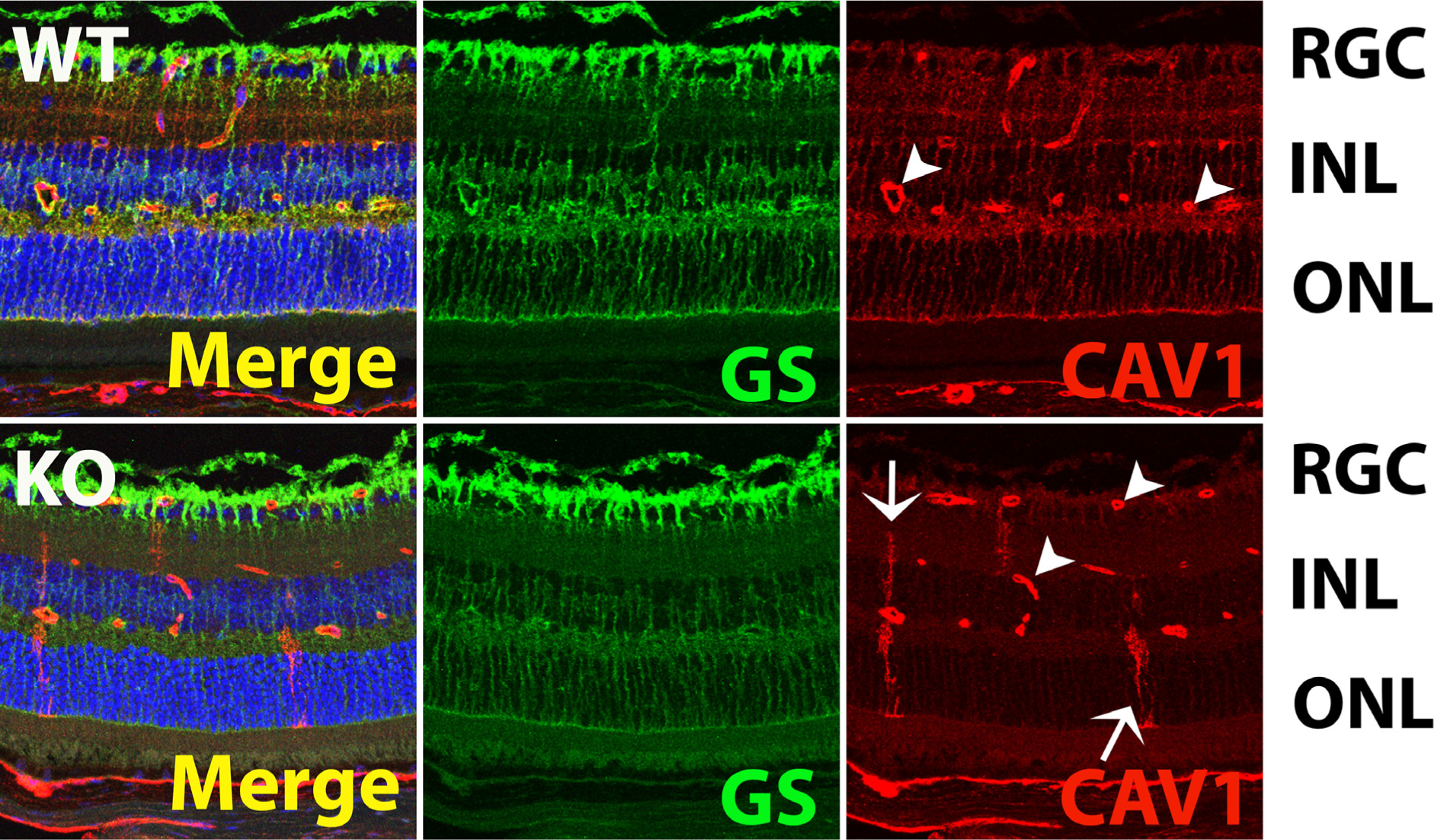
Neuroretinal-specific Cav1 depletion targets retinal Müller glial cells. Immunohistochemical staining of mouse retinal sections stained with anti-CAV1 and costained with Müller cell-specific marker glutamine synthetase (GS) shows high CAV1 protein expression in the retinal Müller glial cell population. Retinal vascular endothelial cross-sections (arrowheads) also exhibit high levels of CAV1 protein expression, but are not targeted by the NR-Cav1 KO model. Sparse mosaic Cre-mediated recombination results in occasional residual Müller glial CAV1 protein expression (arrows). WT, NR-Cav1 WT; KO, NR-Cav1 KO; GS, glutamine synthetase; Cav1, caveolin 1; RGC, retinal ganglion cell layer; INL, inner nuclear layer; ONL, outer nuclear layer.

### Neural retinal Cav1 depletion primarily targets retinal Müller glia CAV1 protein

To ensure that our model specifically targeted Cav1 in the neural retinal compartment, we assessed CAV1 protein expression in retinal tissue sections by immunohistochemistry. In order to deduce retinal cell populations targeted by Chx10-mediated Cav1 depletion, we co-stained retinal tissue sections with cell-specific markers as follows: Müller glia, glutamine synthetase (GS); retinal ganglion cells (RGCs), Brain-Specific Homeobox/POU Domain Protein 3A (Brn3a); bipolar cells (BPCs), Chx10; and rod photoreceptor cells (PRCs), rhodopsin (1D4) (Fig. 2; Fig. S1). We observed the highest levels of CAV1 protein expression in Müller glia cells and retinal blood vessels, which agrees with previous studies (17,22,24,35,36). In retinal tissue sections, Müller glia have a unique morphology where their cellular processes span the retina from inner limiting membrane at the edge of the RGC layer to the outer limiting membrane at the edge of the outer nuclear layer (ONL); whereas, the retinal vasculature appears as discrete vessel cross-sections in the plexiform synaptic layers of the retina (Fig. 2, arrowheads). Importantly, in all sections analyzed, CAV1 expression in the retinal vasculature was not affected by NR-Cav1 depletion (Fig. 2, arrowheads; Fig. S1). However, the characteristic staining of CAV1 in the Müller glia was largely ablated with NR-Cav1 KO, though we did observe modest mosaicism in Müller glia staining (Fig. 2, arrows) as previously reported (34). Alternatively, RGCs of the inner retina (illustrated by Brn3a immunoreactivity) do not express high levels of CAV1 (Fig. S1A). Likewise, BPCs located in the inner nuclear layer (INL) also do not express detectable levels of CAV1 protein (Fig. S1B). Finally, we also did not detect CAV1 immunoreactivity in the ONL region, which houses retinal photoreceptors (Fig. S1C) (37). Thus, our data support that CAV1 protein is most highly expressed in the Müller glia and blood vasculature of the mouse retina. Importantly, Chx10-mediated Cre-recombination of Cav1 significantly reduces Müller glia-specific CAV1 protein while retaining CAV1 protein expression in retinal tissue vasculature. This supports that the majority of CAV1 protein depletion observed in our retinal lysates (Fig. 1) is due to Müller-derived Cav1 recombination. Thus, our NR-Cav1 KO provides an ideal model to assess the contribution of neural retinal Cav1 on the retinal immune response.

### Neuroretinal-Cav1 promotes retinal endotoxin-mediated immune response

We have previously shown that global Cav1 depletion prevented endotoxin-induced elevation of inflammatory retinal cytokines, which suggested that Cav1 functions to *promote* proinflammatory cytokine production. Because we suspected that neural retinal-specific Cav1 expression was responsible for this effect, we used our NR-Cav1 KO model to assess whole retinal cytokine levels following ocular immune challenge via intravitreal LPS injection (Fig. 3). Immune challenge in mice resulted in a robust proinflammatory cytokine response in WT animals (Fig. 3, LPS effect = *). Similar to our previous data collected from global Cav1-KO retinas (21), whole retinal tissue from NR-Cav1 KO animals exhibited lower inflammatory cytokine concentrations (pg/mL) than their WT counterparts following LPS challenge (Fig. 3: IL-6 (interleukin 6): *28.3%*, 165.5 NR-KO vs 585.4 NR-WT; CXCL1/KC (C-X-C motif chemokine ligand 1): *18.1%*, 792.0 NR-KO vs 4374.0 NR-WT; CCL2/MCP-1 (monocyte chemoattractant protein 1): *22.2%* 185.9 NR-KO vs 838.4 NR-WT; and IL-1β (interleukin 1 beta): *71.3%*, 15.9 NR-KO vs 22.3 NR-WT). These data suggest that neural retina-derived Cav1 function is sufficient to promote LPS-induced retinal immune activation. Furthermore, our findings here support that our previous observation of CAV1-dependent cytokine production in global Cav1 KO animals was likely due to CAV1 function in the neural retinal compartment (21).

**Figure 3.**
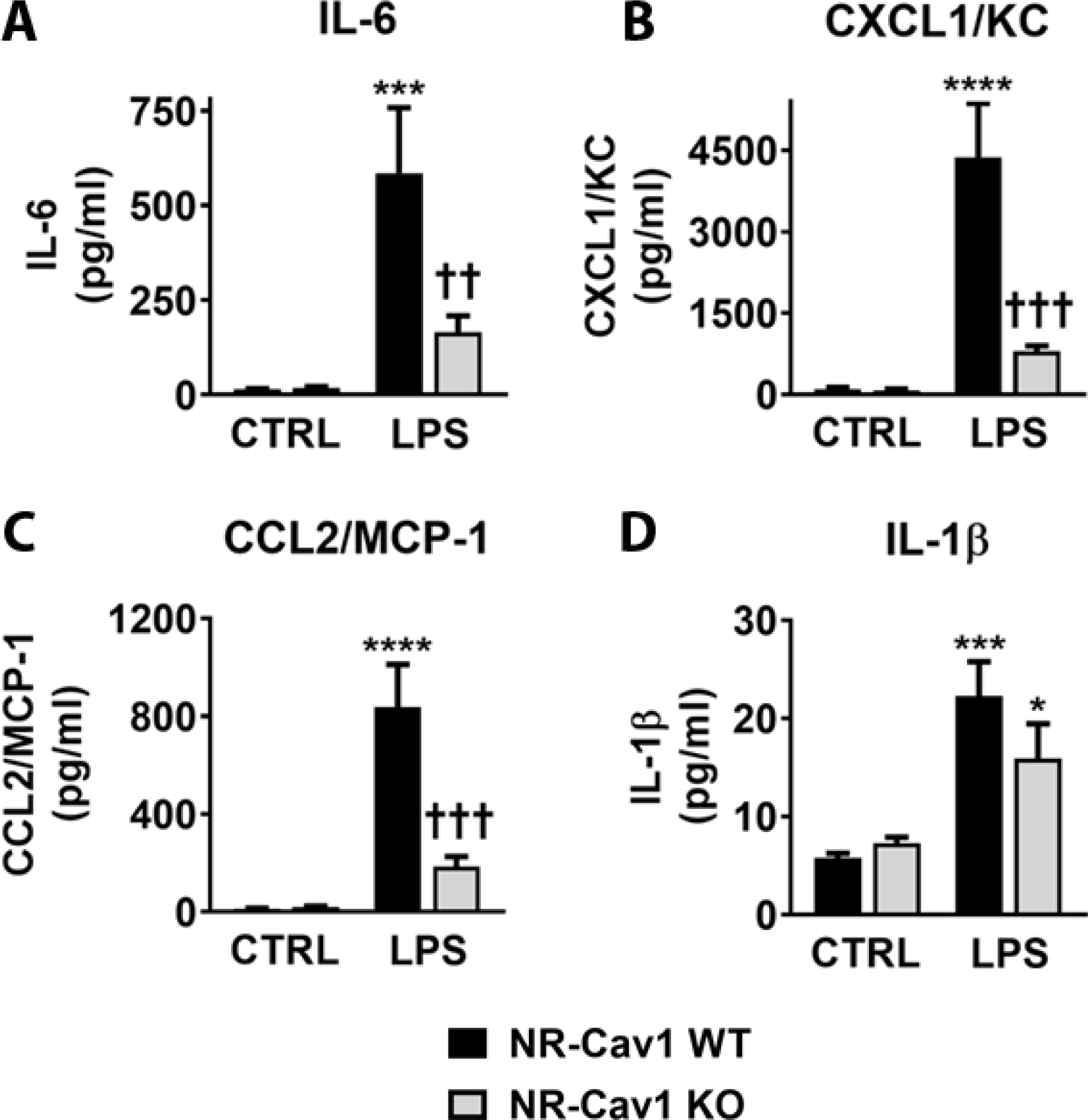
Neuroretinal-specific Cav1 depletion blunts endotoxin-mediated retinal proinflammatory cytokine production. Multiplex protein suspension array analysis showing endotoxin-mediated inflammatory activation in NR-Cav1 WT and KO retinas 24h-post intravitreal LPS injection. Histograms represent measured concentrations of proinflammatory cytokines/chemokines (A) IL-6, (B) CXCL1/KC, (C) CCL2/MCP-1, and (D) IL-1β. Data are mean ± SEM and were analyzed using 2-way ANOVA, with Fisher’s LSD post-hoc. Treatment effect: *<0.05, ***p<0.001, ****p<0.0001; Genotype effect: ††p<0.01, †††p<0.001. IL-6, interleukin 6; CXCL1/KC, C-X-C motif chemokine ligand 1; CCL2/MCP-1, monocyte chemoattractant protein 1; IL-1β, interleukin 1 beta.

Previously, in global Cav1-KO animals, we made the paradoxical observation that retinal immune cell infiltration following LPS stimulation was *elevated* compared to WT mice, which supported that Cav1 functions to *suppress* the retinal immune response. As Cav1 plays a role in several cell processes and is expressed in multiple retinal cell populations (e.g., vascular endothelium), we reasoned that this finding could be explained by differential cell-specific Cav1 functions in the neural retina as opposed to Cav1 function in the endothelia and/or systemic immune system. We, therefore, hypothesized that we would observe *reduced* immune cell infiltration following intraocular LPS challenge when Cav1 depletion is specifically targeted to the neural retina. As suspected, using flow cytometry, we found that NR-Cav1 KO animals exhibited lower levels of retinal immune cell infiltrate than controls, following LPS challenge (Fig. 4A: total leukocytes: *23.2%*, 20,354 NR-KO vs 4,713 NR-WT; PMNs (polymorphonuclear leukocytes): *22.1%*, 4325 NR-KO vs 19,596 NR-WT; INF. MNCs (inflammatory monocytes): *52.9%* 202 NR-KO vs 382 NR-WT; and MΦ (macrophages): *58.7%*, 311 NR-KO vs 530 NR-WT). These data further suggest that neural retinal-specific Cav1 functions to promote the retinal immune response to LPS, and that NR-Cav1 depletion is sufficient to prevent both proinflammatory cytokine production *and* immune cell infiltration into retinal tissue. To further validate that this observation was due to neural retinal compartment-specific effects, we similarly assessed immune infiltration in endothelium-specific Cav1 KO animals (*Tie2*-Cre Cav1 KO model; Endo-Cav1 KO). Endo-Cav1 KO mice did <u>*not*</u> exhibit the suppressive effect on infiltration, which further supports that neural retinal Cav1, specifically, plays a crucial role in retinal immune activation (Fig. 4B: total leukocytes: 2,741 NR-KO vs 2,435 NR-WT; PMNs: 2,164 NR-KO vs 1,799 NR-WT;INF. MNCs: *163* NR-KO vs 184 NR-WT; and MΦ: 445 NR-KO vs 441 NR-WT).

**Figure 4.**
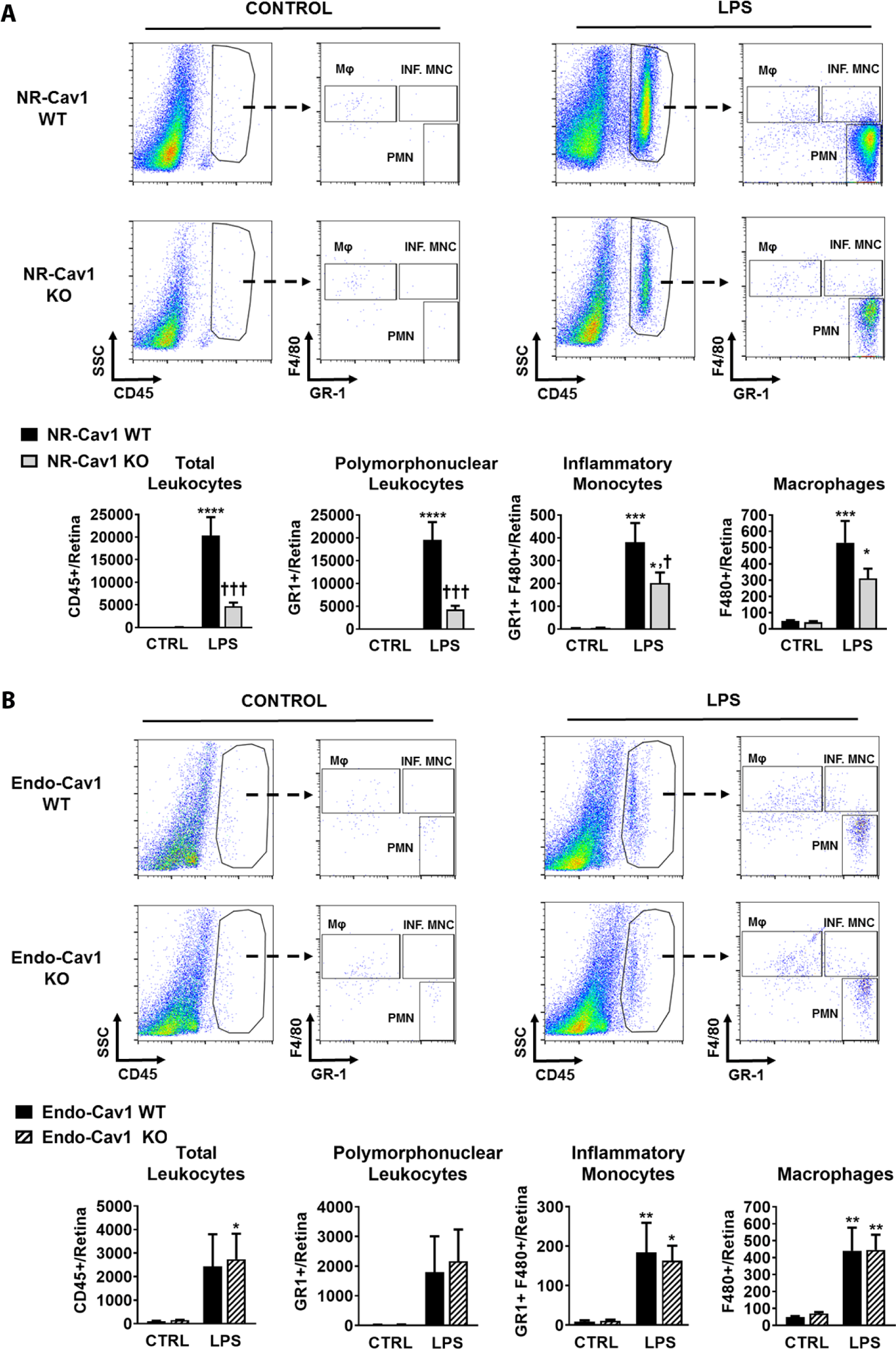
Neuroretinal-specific (but <u>not</u> endothelial cell-specific) Cav1 depletion reduces immune cell recruitment to retinal tissue. Flow cytometry data representing immune cell infiltration into retinal tissue 24-hours post-intraocular LPS challenge. Animals were perfused prior to collection of whole retinal tissue from Chx10-Cav1 WT and Chx10-Cav1 KO animals. (A) Representative dot plots showing spread of all data and gating strategy represent the following cell populations: total leukocytes (CD45^hi^), PMNs (F4/80^−^/GR1^+^), INF. MNCs (F4/80^+^/Gr1^+^), and MΦ (F4/80^+^/GR1^−^). (B) Histograms of summarized data from control or LPS-treated Chx10-Cav1 WT and Chx10-Cav1 KO animals. Data are mean ± SEM and were analyzed using 2-way ANOVA, with Fisher’s LSD post-hoc. Treatment effect: *p<0.05, **p<0.01, ***p<0.001, ****p<0.0001; Genotype effect: †p<0.05, †††p<0.001. PMNs, polymorphonuclear leukocytes; INF. MNCs, inflammatory monocytes; MΦ, macrophages.

### Immunomodulatory TRAF3 protein is highly expressed in retinal tissue and is upregulated in NR-Cav1 KO animals that exhibit reduced immune responses

While this work is focused on clarifying the role of NR-specific Cav1 function, we ultimately hope to identify novel potential regulators of retinal inflammation that may be targeted in order to prevent retinal damage. Here, our work suggests that NR-Cav1 *promotes* retinal inflammation and that an active immunosuppressive mechanism is present in the absence of NR-Cav1 expression. Thus, we have begun to investigate potential immune suppressors that are *upregulated* in response to Cav1 depletion.

CAV1 protein is primarily localized to the cell membrane where it participates in formation and stabilization of specialized lipid raft domains, which can facilitate regulatory effects on membrane receptor localization and activation (38). Thus, because we suspected that Cav1 regulates immune pathways at the level of membrane receptor complex formation and signal activation, we decided to narrow our search for novel immune regulators to those that interact with CAV1 protein at the cell surface. Thus, we isolated retinal membrane fractions (CAV1-containing membrane fractions) from whole retinal tissue and performed mass spectrometry and quantitative proteomics analysis on membrane-associated proteins using previously described methods (22). Our data show significant downregulation of 108 genes (including Cav1) and upregulation of 177 genes in retinal membrane fractions in response to NR-Cav1 deletion. Data for the top 70 differentially expressed proteins are represented in Fig. 5 (blue = downregulated; red = upregulated). Retinas from our NR-Cav1 KO animals exhibited 70% reduction in CAV1 peptide abundance (Fig. 5), which is consistent with the 76% reduction observed by Western blotting of whole retinal tissue lysates (Fig. 1). Collectively, our data suggest that the majority of retinal CAV1 protein resides within the plasma membrane of neural retinal cell populations (i.e., primarily the Müller glia; Figs. 1,2,5). After validating that NR-Cav1 KO results in CAV1 protein reduction in retinal membrane fractions, we then used the ShinyGO gene ontology enrichment analysis online software to identify which of our 177 upregulated genes are functionally associated with immune and stress response pathways (Table 1).

**Table 1.** Effect of neuroretinal-specific depletion on immune-related gene ontology categories for significantly upregulated genes determined by proteomics data. ShinyGO analysis was performed on all 177 genes that were identified by proteomics as being significantly upregulated following NR-Cav1 depletion. Table represents immune-associated functional category groups (“High level GO category”) defined by high-level gene ontology terms used by ShinyGO analysis software. Table also includes both the number (“N”) and gene names (“Gene”) of upregulated genes identified to contribute to the immune-associated GO categories.

| Immune-Related Gene Ontology Enrichment Analysis for All 177 Significantly Upregulated Genes following NR-Cav1 Depletion |  |  |
| --- | --- | --- |
| N | High level GO category | Genes |
| 45 | Response to stress | DUSP3 RBM14 ANXA6 RAD21 HSPA9 ANXA5 VCP PCK2 H2AFX UBE2N ENTPD2 VAMP4 PPP3CA GSTM5 GORASP2 UCHL5 XAB2 <u>TRAF3</u> OXCT1 DLG1 FEN1 CORO1B HSPD1 CAPN2 DDAH1 SFPQ NONO NEDD4 ADRA2A MACROD1 NPLOC4 SMC1A CORO2B HNRNPA1 USP14 PRKCA PRMT1 ANXA7 UCHL1 PTPN11 WDFY1 GSTM7 CAMK2G HSPA4L XRN2 |
| 17 | Immune system process | DUSP3 RBM14 HSPD1 VAMP4 <u>TRAF3</u> HSPA9 SFPQ NONO NPLOC4 USP14 RPS17 PRMT1 PTPN11 DLG1 NEDD4 WDFY1 UBE2N |
| 13 | Immune response | DUSP3 RBM14 VAMP4 <u>TRAF3</u> SFPQ NONO NPLOC4 USP14 DLG1 NEDD4 WDFY1 HSPD1 UBE2N |
| 12 | Regulation of immune system process | DUSP3 RBM14 HSPD1 <u>TRAF3</u> HSPA9 SFPQ NONO NPLOC4 PRMT1 DLG1 WDFY1 UBE2N |
| 9 | Activation of immune response | DUSP3 RBM14 SFPQ NONO NPLOC4 <u>TRAF3</u> WDFY1 HSPD1 UBE2N |
| 4 | Immune effector process | <u>TRAF3</u> NPLOC4 DLG1 HSPD1 |

**Figure 5.**
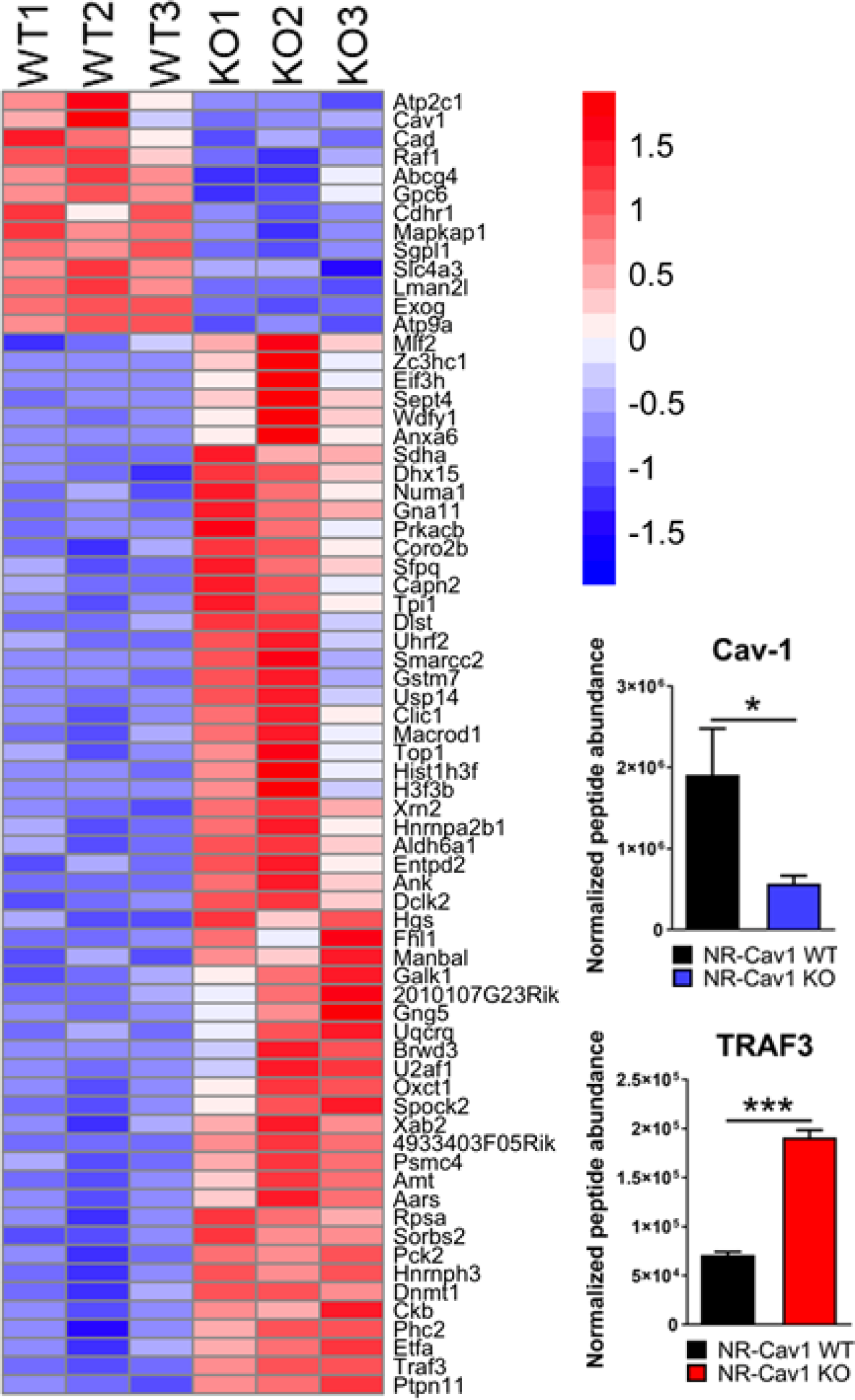
Effect of neuroretinal-specific depletion on membrane-associated protein composition. Mass spectrometry-based proteomics heat map analysis (*pheatmap* R) of membrane raft fractions from whole retinal tissue and histograms representing peptide abundance quantification that shows Cav1 downregulation and TRAF3 upregulation in raft protein fractions as a result of NR-Cav1 depletion. Each replicate sample contained pooled retinas from 8 mice. Blue = downregulated; red = upregulated. WT, NR-Cav1 WT; KO, NR-Cav1 KO. Data are mean ± SEM and were analyzed via Student’s t-test (* p<0.05; *** p<0.001). WT, NR-Cav1 WT; KO, NR-Cav1 KO.

Interestingly, we identified upregulation of membrane TRAF3 in NR-Cav1 retinas, which we validated with retinal membrane fractionation and TRAF3 Western blot analysis (Fig. 5; Fig. S2). Traf3 is a known inflammatory regulator in non-retinal tissues that associates with immune receptor complexes at the cell membrane in order to modulate downstream signaling (39–42). Importantly, Traf3 function in the retina has not yet been investigated, though existing proteomics data sets suggest that Traf3 is highly expressed in the retina compared to other tissues (Genecards). This, together with our current data, suggests that Traf3 likely functions to regulate immune responses in the retina, which we intend to examine in future studies.

To better understand how CAV1 might interact with TRAF3 to modulate retinal immune responses, we used the STRING v10 database to obtain protein-protein interaction (PPI) information for the most significantly changed proteins (adjusted p-value <=0.10) within our data set (32). Fig. 6 shows that CAV1 is known to share at least 19 PPIs with 7 proteins and illustrates how loss of Cav1 might impact the activity and/or upregulation of 4 proteins (including TRAF3) (Fig. 6; Fig. S3). At least 3 of the 4 CAV1-TRAF3 shared interacting proteins have been associated with TNF-alpha-induced cell death (i.e., RIPK1/Receptor-interacting serine/threonine-protein kinase 1, TRAF2/ tumor necrosis factor (TNF) receptor-associated factor 2, TRADD/ tumor necrosis factor receptor type 1-associated DEATH domain protein) (43,44). The bulk of the PPIs shared with CAV1 are among TRAF3 and PTPN11 (tyrosine-protein phosphatase non-receptor type 11), which TRAF3 is known to inhibit. Many of these shared PPIs (e.g., EGF/ epidermal growth factor, SRC/neuronal proto-oncogene tyrosine-protein kinase, BCAR1/breast cancer anti-estrogen resistance protein 1) are also associated with anoikis (programmed cell death upon detachment from the extracellular matrix). Indeed, CAV1 has been reported to increase resistance to anoikis, which may be related to Cav1’s role in neuroprotection (45,46). But because the changed proteins were not significantly enriched in immune functions or pathways, we cannot rule out the possibility that the immune effects of Cav1 KO may be indirect and due to alterations in the structure of the plasma membrane. This may, in fact, help reconcile why some studies observe a pro- or anti-inflammatory effect, as a change in membrane dynamics would likely affect both.

**Figure 6.**
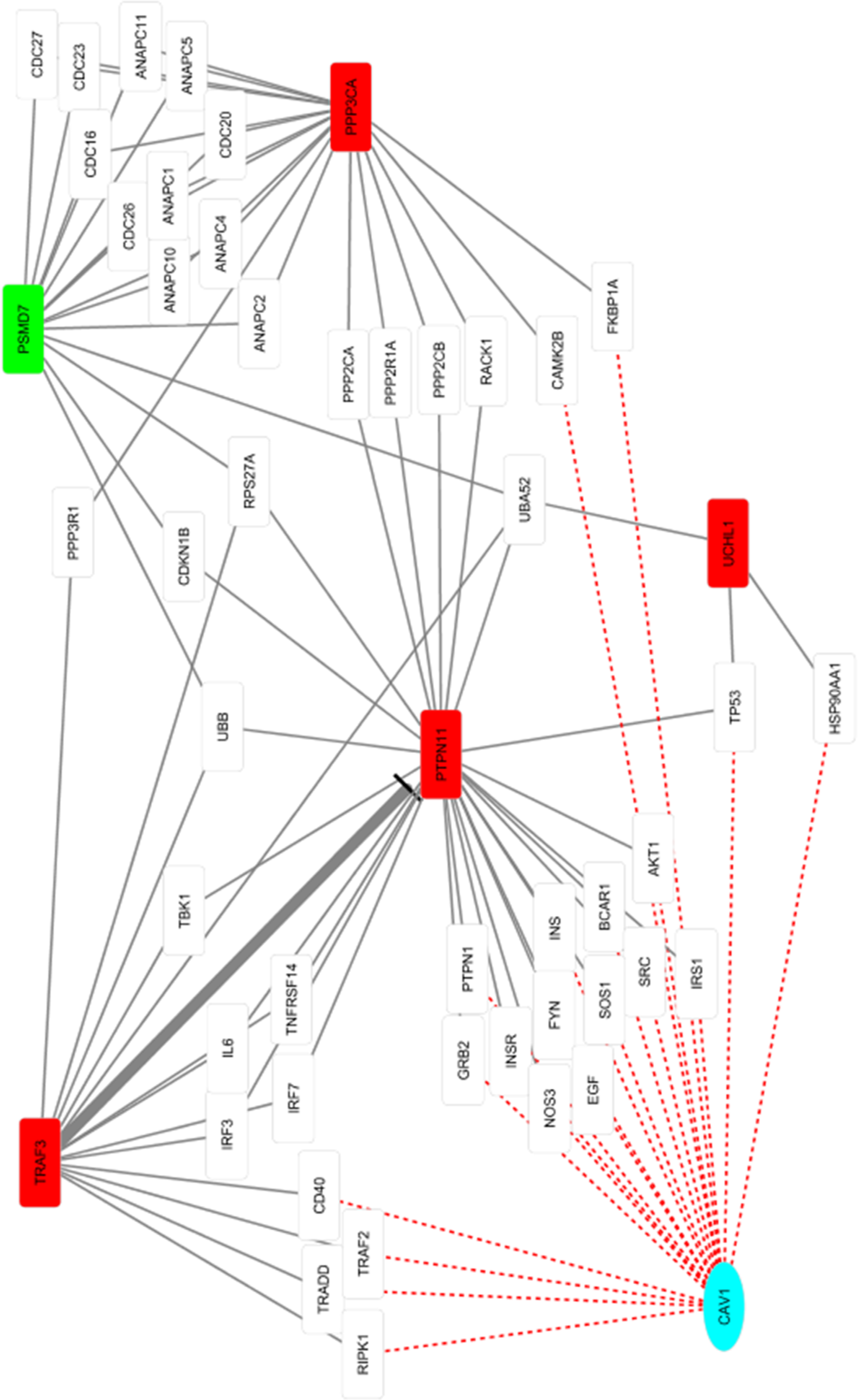
CAV1 protein interaction network. Shared protein-protein interactions (PPIs) between CAV1 and the most significantly changed proteins. Red=upregulated after CAV1 KO, green=downregulated. Red dashed lines indicate potential PPIs that are altered by an absence of cellular CAV1 protein.

## Discussion

Previous work from our lab showed that Cav1 depletion in global-Cav1 KO animals prevented inflammatory cytokine production in the retina, which suggested that Cav1 promotes inflammation in retinal tissue (21). Our current results determine that NR-Cav1 depletion is sufficient to produce this response. Likewise, this work shows that NR-Cav1 depletion also prevents immune cell infiltration. However, our previous results illustrate that global depletion of Cav1 actually *exacerbates* retinal immune cell infiltration. Together, this suggests that ablation of a non-neural retinal Cav1 pool is responsible for the enhanced presence of immune cells in the retinal tissue of global-Cav1 KO animals. We initially suspected that the elevated immune cell infiltration in the global-Cav1 KO animals could be explained by enhanced vascular permeability due to endothelial-Cav1 depletion. Thus, we also assessed infiltration in our endothelial (*Tie2*)-Cav1 KO animals. Interestingly, there was no difference in infiltration values between Endo-Cav1 KO animals compared to WT controls. Thus, endothelial-derived Cav1 is *not* responsible for the effect of elevated infiltration observed in the global-Cav1 KOs. We believe, rather, that immune cell-specific effects of Cav1 depletion in global-Cav1 KO animals is responsible. Previous work from our lab has shown that global-Cav1 KO animals have more CD45-positive cells (i.e., leukocytes) within their retinal vasculature compared to WT animals (21). Thus, we suspect that the increased presence of circulating immune cells in global-Cav1 KO animals results in greater accessibility to the retinal vessels under basal conditions and primes the retina to uptake more immune cells following an immune response that induces vessel permeability. Alternatively, the enhanced infiltration phenotype observed in the global-Cav1 KO could be explained by loss of RPE-specific Cav1-dependent functions, which are also not targeted in our NR-Cav1 KO animals.

Our data support that the majority of neural retinal (and total retinal) Cav1 protein resides within the Müller glial cell population, which is in agreement with ours and other previous work (24). In addition to their important structural and neurosupportive roles in the retina (i.e., metabolic homeostasis, neurotransmitter recycling, and intercellular communication), Müller glia also play an unappreciated role in retinal immunity (47). Müller glia have been shown to express innate immune receptors and to produce inflammatory and neuroprotective cytokines (48,49). Intriguingly, Cav1 has been shown to be important for secretion of cytokines such as IL-6 (50,51). Furthermore, previous data from our lab suggests a Cav1-dependent role in IL-6 family cytokine production that is upstream of neuroprotective signaling (via STAT3 pathway activation) (24). These studies are consistent with our observed suppression of LPS-induced cytokine production in NR-Cav1 KO mice and support a possible role for Müller-derived inflammatory cytokine production and secretion following ocular insult. However, further studies are needed to fully understand the contribution of Müller glia-derived immune responses and the role of Müller glia in maintenance of the retinal immune-privileged environment.

Many studies have been published on Cav1 since its identification in the late 1980s. However, the precise role of Cav1 in immune regulation has remained unclear. Furthermore, whether Cav1 generally functions to promote or suppress retinal immune activation is equally as perplexing. Our study suggests that neural retinal Cav1, specifically, *promotes* retinal inflammation. Because Cav1 is known to stabilize cell surface receptors in specialized lipid raft membrane domains (38), it is conceivable that the blunted inflammatory activation observed with NR-Cav1 depletion is due to altered receptor-membrane dynamics that affect immune receptor complex formation and propagation of downstream inflammatory signaling pathways. Thus, we have begun to investigate novel potential inflammatory regulators that might associate with the cell membrane to regulate receptor activation in the absence of Cav1. Interestingly, we identified elevated TRAF3 protein levels in retinal membrane fractions from our NR-Cav1 KO animals. While Traf3 is known to play an important immune regulatory role in other tissues, the function in the retina is unknown. Additionally, Traf3 has been shown to directly interact with immune receptors, including CD40 and TLR4, which are both expressed in Müller glial cells (52). Thus, we suggest that Traf3 may play a previously-unappreciated role in retinal inflammatory modulation. Our future studies include investigation of Traf3 function in the retina, as well as further examination into the mechanism(s) of Cav1-dependent inflammatory activation in Müller glia.

## Experimental procedures

### Animals

NR-specific Cav1 KO mice (Chx10-Cre; Cav1^flox/flox^) were generated using Cre-Lox technology (24). Animals carrying Chx10-promoter-driven Cre recombinase (stock#: 005105, The Jackson Laboratory, Bar Harbor, ME) were bred with animals carrying the Cav1 gene flanked by Lox P sites located upstream and downstream of exon 2 (25). Endothelial-specific Cav1 knock-out mice (Tie2-Cre; Cav1^flox/flox^) were generated similarly, using endothelial cell-specific recombination (B6.Cg-Tg (Tek-cre) 1Ywa/J; stock#: 008863, The Jackson Laboratory, Bar Harbor, ME) (26). Cre-negative littermates were used as controls. Mice were screened for rd1 and rd8 mutations prior to this study. All procedures were approved by the University of Oklahoma Health Sciences Center (OUHSC) Institutional Animal Care and Use Committee (IACUC) and comply with the Association for Research in Vision and Ophthalmology (ARVO) Statement for the Use of Animals in Ophthalmic and Visual Research.

### Whole retinal tissue lysis

Total retinal protein was obtained via whole retinal tissue lysis by brief sonication in 2X lysis buffer (120 mM octyl glucoside, 2% Triton X-100, 20 mM Tris-HCl, pH 7.4, 200 mM NaCl, 1 mM EDTA) containing a protease inhibitor cocktail (Roche). Lysates were cleared by centrifugation at 13K rpm, at 4°C. Protein concentration was determined via BCA assay (ThermoSci, Cat# 23225) and samples were diluted in 6X Laemmli buffer.

### Western blotting

Retinas were homogenized in buffer containing 100 mM sodium carbonate and 1 mM EDTA using an plastic pellet pestle and brief pulse sonication on ice. Membranes were isolated by centrifugation at 16,000 x g in a benchtop microcentrifuge and membrane pellets were lysed in 1X lysis buffer. Protein content of lysates was determined by the BCA assay with bovine serum albumin as the standard. Protein separation was achieved via SDS-PAGE and proteins were transferred to nitrocellulose for immunoblotting with the following primary antibody dilutions: rabbit monoclonal anti–Caveolin-1 (1:1000; Cell Signaling #3267), mouse monoclonal anti–Actin (1:1000; ThermoFisher #MA1-744), rabbit polyclonal anti-TRAF3 (TNF Receptor-Associated Factor 3; 1:500; Novus Biologicals, #NBP-88639), and rabbit polyclonal anti-Gαt (Transducin; 1:4000; Santa Cruz). Immunoreactivity was detected using species-appropriate horseradish peroxidase (HRP)-conjugated secondary antibodies (1:5000; GE Healthcare, Cleveland, OH, USA). Western blots were imaged via HRP chemiluminescent detection (Azure Biosystems, Inc.; Dublin, CA) and densitometric analyses were performed using Image Studio Lite software (LI-COR Biosciences).

### Immunohistochemistry & confocal microscopy

Whole eye globes from euthanized adult mice were fixed in Prefer fixative (Anatech, Ltd., Battlefield, MI), embedded in paraffin, and cut into 5-μm sections. Sections were deparaffinized, permeabilized with 1% Triton X-100 in PBS, then incubated with blocking solution (10% normal horse serum, 0.1% Triton X-100, in PBS) for one hour prior to immunohistochemistry, which was performed as previously described (27,28) using the following antibodies: rabbit monoclonal anti-Caveolin-1 (1:100, Cell Signaling #3267); mouse monoclonal anti-Glutamine Synthetase (for Müller glia-specific marker; 1:5000 Millipore #MAB302, clone GS-6); goat polyclonal anti-Brn3a (for retinal ganglion cell marker, 1:100, Santa Cruz #sc-31984); sheep polyclonal anti-Chx10 (for bipolar cell marker; 1:100, Exalpha #X1179P); and anti-rhodopsin/1D4 (for photoreceptor marker; 1:100, gifted from R. Molday, Univ. of British Columbia). Immunoreactivity was detected with Alexa Fluor-conjugated secondary antibodies (1:500, Life Technologies, Grand Island, NY) and nuclei were stained with DAPI (4,6-diamidino-2-phenylindole). Images of immunostained retinal tissue sections were acquired on an Olympus FV1200 laser scanning confocal system using FluoView software (Olympus). Pseudocolors were assigned to images as follows: Glutamine Synthetase, Brn3a, Chx10, or 1D4 (green); Cav-1 (red); and nuclei (blue).

### Endotoxin-induced uveitis model

As previously described, local, intravitreal injection of 1 μg LPS (*Salmonella typhimurium*; Sigma) was used to induce ocular inflammation (21,29). Mice were anesthetized via intraperitoneal injection of ketamine (100 mg/kg)/xylazine (10 mg/kg). Each eye was then injected with either 1 μL LPS diluted in sterile PBS vehicle or PBS alone using a 10 µL glass syringe (Hamilton).

### Cytokine/chemokine quantification Assay

Retinas from euthanized animals were collected 24 h after LPS or PBS injection. Retinal tissue dissection was performed using a stereo dissecting microscope (Zeiss Stemi DV4; Carl Zeiss Microscopy, Jena, Germany). Retinas were processed and cytokines protein levels were measured by BioPlex suspension array (Bio-Rad Life Science, Hercules, CA, USA) as previously described (21).

### Flow cytometry

Flow cytometry was performed on retinal samples as described previously for neural tissues with minor modification (21,30). Twenty-four hours after LPS or PBS injection, mice were perfused with PBS, and single-cell suspensions were obtained from retinas by digestion of minced tissue with liberase TL in RPMI 1640 at 37°C for 30 minutes. Next, the cell suspensions were passed through a 40-µm nylon cell strainer (Thermo Fisher Scientific) followed by washing with staining buffer (SB; PBS supplemented with 2% FBS, and 2 mM EDTA). Samples were blocked with anti-mouse CD16/32 (Fc-block, Invitrogen, San Jose, CA, USA) for 10 minutes at 4°C followed by staining with antibody cocktail containing Pacific-Blue-conjugated anti-mouse CD45 (clone 30-F11; Biolegend, San Diego, CA, USA), PE-conjugated anti-mouse Ly-6G (Gr-1) (clone RB6-8C5; Biolgend), and APC-conjugated anti-mouse F4/80 (clone BM8; Biolegend). Samples were incubated 20 minutes at 4°C and then washed with SB buffer. The data were acquired immediately after staining using a MacsQuant flow cytometer (Miltenyi Biotec, Auburn, CA USA) and analyzed with FlowJo software (FlowJo LLC). The events were gated by forward and side scatter, as well as CD45 to distinguish infiltrating immune cells (CD45^hi^).

### Mass spectrometry-based proteomics

Membrane proteins were prepared from whole frozen retinal tissue as described previously (22). Briefly, retinal tissues were lysed in high salt buffer (2M NaCl, 10mM HEPES-NaOH, pH 7.4, 1mM EDTA, complete protease inhibitors). Pellets were harvested via high-speed centrifugation and were subsequently homogenized in carbonate buffer (0.1M Na_2_CO_3_, pH 11.3, 1mM EDTA, complete protease inhibitors). Pellets were collected and re-extracted with carbonate buffer a second time. Pellets were collected and re-homogenized in urea buffer (4 M urea, 100 mM NaCl, 10 mM HEPES/NaOH, PH 7.4, 1 mM EDTA, complete protease inhibitors). The resulting membrane fractions were collected via centrifugation and were used for tryptic digest as described previously (22). Label-free mass spectrometry was used to measure peptide abundances from membrane-enriched fractions of retinal samples from both NR-Cav1 WT and NR-Cav1 KO mice. For an in-depth description of the LC-MSMS procedure and detailed label-free peptide quantification analysis parameters, please refer to Hauck, et al., 2010. Briefly, peptide spectra obtained by LC-MSMS on an Ultimate3000 nano HPLC system (Dionex, Sunnyvale, CA) online coupled to a LTQ OrbitrapXL mass spectrometer (Thermo Fisher Scientific) were transformed to peak lists using Progenesis software, followed by label-free quantification based on peak intensities. Total list of identified peptides and proteins are available in *Supporting Information* (Supplemental Table 1). The mass spectrometry data have been deposited to the ProteomeXchange Consortium via the PRIDE (31) partner repository with the dataset identifier PXD016872 and 10.6019/PXD016872. Reviewer account details:

**Password:** VbXalKFX

### Bioinformatics

Peptide abundances were log2-transformed and analyzed for differential abundance using the *limma* R package. The row-scaled abundances of the top 70 differentially expressed retinal proteins affected by NR-Cav1 depletion were visualized as a heatmap using the *pheatmap* R package (blue = downregulated; red = upregulated). The resulting complete proteomics data set was analyzed using both pathway and network analyses. KEGG (Kyoto Encyclopedia of Genes and Genomes) analysis and IPA (Ingenuity Pathway Analysis) were used for pathway enrichment in order to identify known gene sets affected by NR-Cav1 depletion. We obtained the protein-protein interaction (PPI) information for the most significantly changed proteins (adjusted p-value <=0.10) from the STRING v10 database (32). Single PPIs (nodes with only one edge) were removed to reduce clutter and to focus upon *shared* PPIs within this network, and visualized with Cytoscape 3.7 (33).

## Acknowledgments

Data are available via ProteomeXchange with identifier PXD016872.

We thank Linda Boone in the NEI P30-supported Cellular Imaging Core for expert preparation of specimens and tissue sections for immunohistochemistry. We are also grateful to Renee Selleck from the Carr laboratory for preparation of retinal tissues for flow cytometry.

## FOOTNOTES

This work was funded by grants from R01EY019494 to MHE; T32EY023202; P30EY021725; an unrestricted grant from Research to Prevent Blindness to the Department of Ophthalmology at the Dean McGee Eye Institute (OUHSC); the German Research Foundation (DFG: HAU 6014/5-1, priority program SPP 2127) to SMH.

Dr. Jami Gurley is the recipient of the Postdoctoral OCAST Fellowship grant HF18-008.

## The abbreviations used are

Cav-1: caveolin-1
NR: neural retina
LPS: lipopolysaccharide
Traf3: TNF receptor-associated factor 3
TNF: tumor necrosis factor
TLR4: toll-like receptor 4
RPE: retinal pigment epithelium
KO: knockout
WT: wild type
GS: glutamine synthetase
RGCs: retinal ganglion cells
Brn3a: Brain-Specific Homeobox/POU Domain Protein 3A
BPCs: bipolar cells
PRCs: photoreceptors
ONL: outer nuclear layer
INL: inner nuclear layer
IL-6: interleukin 6
CXCL1: C-X-C motif chemokine ligand 1
MCP-1: monocyte chemoattractant protein 1
PMNs: polymorphonuclear leukocytes
INF. MNCs: inflammatory monocytes
MΦ: macrophages
PPI: protein-protein interaction
RIPK1: receptor-interacting serine/threonine-protein kinase 1
TRAF2: TNF receptor-associated factor 3
TRADD: tumor necrosis factor receptor type 1-associated DEATH domain protein
PTPN11: tyrosine-protein phosphatase non-receptor type 11
SRC: neuronal proto-oncogene tyrosine-protein kinase
BCAR1: breast cancer anti-estrogen resistance protein 1
DAPI: 4,6-diamidino-2-phenylindole
KEGG: Kyoto Encyclopedia of Genes and Genomes
IPA: Ingenuity Pathway Analysis.

